# Chloroflexi persisting for millions of years in oxic and anoxic deep-sea clay

**DOI:** 10.1101/2020.05.26.116590

**Authors:** Aurèle Vuillemin, Zak Kerrigan, Steven D’Hondt, William D. Orsi

## Abstract

Chloroflexi are widespread in energy-limited subseafloor sediments, but how Chloroflexi respond to subseafloor energy limitation under oxic and anoxic conditions is poorly understood. Here, we characterize the diversity, abundance, activity, and metabolic potential of Chloroflexi in oxic and anoxic abyssal clay from three deep-sea cores covering up to 15 million years of sediment deposition, where Chloroflexi are a major component of the community throughout the entire cored sequence at all sites. In oxic red clay at two different sites, Chloroflexi communities exhibit net death over both 10-15 million year cored sequences, and gene expression was below detection despite the availability of oxygen as a high energy electron acceptor, indicating a reduced level of activity. In contrast at the anoxic site, Chloroflexi abundance and gene expression increase below the seafloor and peak in 2 to 3 million year old sediment. The anaerobic subseafloor Chloroflexi exhibited a homoacetogenic metabolism and potential for energetically efficient intracellular H_2_ recycling that have been proposed to confer a fitness advantage in energy-limited subseafloor habitats. Our findings indicate that the expression of this energy efficient metabolism in Chloroflexi coincides with net growth over million year timescales in deep-sea anoxic clay.

**Significance statement:** Chloroflexi are widespread in energy-limited subseafloor sediments, both in oxic subseafloor sediments that are energetically limited by the availability of electron donors (organic matter) and in anoxic sediments that are energetically limited by the availability of high energy terminal electron acceptors. How Chloroflexi respond to these different forms of energy limitation over long time scales is poorly understood. We present new data that demonstrates how key differences in metabolism are manifested in different communities of aerobic and anaerobic Chloroflexi subsisting over millions of years in oxic and anoxic deep-sea clay. These data provide new insights into how certain Chloroflexi respond to different types of long-term energy limitation.

## Introduction

Vast regions of the dark ocean have extremely slow sedimentation rates, ranging from 1 to 5 meters of sediment deposition per million years in abyssal regions (D’Hondt et al., 2015) and most organic flux to the sediment is consumed before it can be buried (Røy et al., 2012). As a result, microbial communities in abyssal subseafloor sediment are characterized by extremely low cell densities (Kallmeyer et al., 2012), experience extreme energy limitation over the long term (Hoehler and Jørgensen, 2013), even over million year timescales under both oxic and anoxic conditions (D’Hondt et al., 2015). The sedimentary material of the deep-sea is characterized by ultra-small (<0.2 µm) clay particles that are highly pressurized under the >5,000 m of overlying seawater, which likely hinders the motility of cells in this extreme environment, resulting in a complete separation from the surface after deposition.

Cells subsisting in deep-sea clays experience one of two types of energy limitation, limitation by electron donors or limitation by electron acceptors. In anoxic deep-sea clay, high energy electron acceptors of oxygen and nitrate are consumed at the sediment surface, and a main terminal electron acceptor is sulfate (Parks et al., 2005; Jørgensen et al., 2019). In contrast, the cells subsisting in subseafloor oxic deep-sea clay are constantly exposed to high energy electron acceptors of oxygen and nitrate throughout the entire sediment column (Røy et al., 2012), but are rather limited by the availability of electron donors in the form of organic matter (D’Hondt et al., 2015). This happens because the sedimentation rates are so slow in the oligotrophic ocean, that what little organic matter makes it to the seafloor is exposed to oxic conditions for exceptionally long timescales before it is buried, making the total concentration of subseafloor organic matter in oxic deep-sea red clay extremely low (Bradley et al., 2019). The end result is an oxic microbial subseafloor community that is limited by electron donors, compared to anoxic communities that have more organic matter but are energetically limited by electron acceptors (Orsi, 2018). The composition of these communities is shaped by selective survival of taxa pre-adapted to these conditions (Starnawski et al., 2017; Kirkpatrick et al., 2019) and includes many poorly characterized clades of Chloroflexi (Inagaki et al., 2006; Nunoura et al., 2018). However, the structure of these communities, their relationship to redox stratification, and their modes of selection as main constituents of the deep biosphere remain poorly constrained from the water-sediment interface down through the deep subseafloor sediment column (Durbin and Teske, 2011; Walsh et al., 2016; Petro et al., 2019).

Chloroflexi are environmentally widespread. Their biogeography and modes of adaptation have been examined in various settings (Biddle et al., 2012) with variable salinities (Mehrshad et al., 2018), including shallow marine (Wilms et al., 2006; Petro et al., 2019), deep lacustrine (Vuillemin et al., 2018a; Kadnikov et al., 2019) and aquifer sediments (Hug et al., 2013) where they have been found to be abundant and highly diverse. They have also been reported as a predominant phylum in sewers (Krumins et al., 2018), anaerobic wastewater treatment systems (Shu et al., 2015), methanogenic reactors (Bovio et al., 2018), marine sponge microbiomes (Bayer et al., 2018) and deep subseafloor sediments (Orsi, 2018). Considering the scale of the marine subsurface (Kallmeyer et al., 2012), subseafloor Chloroflexi may be one of the most abundant microbial components on Earth and may drive key biogeochemical cycles on a global scale (Zinger et al., 2011; Hug et al., 2013; Fincker et al., 2020). However, many groups of subsurface microorganisms, including the Chloroflexi, remain uncultivated (Biddle et al., 2012; Solden et al., 2016). Thus, their metabolic properties remain poorly characterized, and the reasons for their broad ecological distribution and diversity have been elusive (Kaster et al., 2014; Wasmund et al., 2014; Sewell et al., 2017).

Within representative isolates among classes of Anaerolineae and Dehalococcoidia, diverse metabolic strategies include anoxic phototrophy (Klatt et al., 2013), fermentation (Yamada et al., 2006, 2017), and reductive dehalogenation (Duhamel and Edwards, 2006; Matturo et al., 2017). In terms of carbon cycling in marine sediments, the transformation of halogenated compounds has been studied extensively (Kittelmann and Friedrich, 2008; Kaster et al., 2014; Atashgashi et al., 2018; Kawai et al., 2014), whereas metagenome-assembled genomes (MAGs) and single-cell amplified genomes (SAGs) provide parallel evidence for respiration of sugars, fermentation, CO_2_ fixation via the Wood-Ljungdahl (W-L) pathway and acetogenesis with substrate-level phosphorylation (Hug et al., 2013; Sewell et al., 2017). The presence of genes encoding dissimilatory sulfite reductase (*Dsr*) and reversible adenylylsulfate reductase (*apr*) imply roles in sulfur cycling and respiration (Wasmund et al., 2017; Mehrshad et al., 2018; Vuillemin et al., 2018b). In addition, the phylum Chloroflexi exhibits a variety of complex monodermic cell envelope structures (Sutcliffe, 2011) and carries genes for multiple environmental adaptations across clades. These adaptations include protection against oxygen and osmotic stress (Wasmund et al., 2014) and aerobic oxidation of H_2_, CO and sulfur compounds at sub-atmospheric concentrations (Islam et al., 2019). Altogether, these metagenomic data suggest versatile respiratory modes with intricate heterotrophic and lithoautotrophic metabolisms, which may support long-term survival in dormant states and dispersal in deep marine environments (Jørgensen and Marshall, 2016; Fullerton and Moyer, 2016).

In previous studies of slowly accumulating oxic and anoxic subseafloor clay from the abyssal North Atlantic covering up to 15 million years of depositional history (Vuillemin et al., 2018c and 2019), we found that Chloroflexi were a major component of the community in both settings. This abundance across multiple coring locations with varying redox states (oxic vs. anoxic) prompted a focused investigation of how the associated differences in long-term energy limitation (i.e. electron donors at oxic sites vs. electron acceptors at anoxic sites) select for metabolic features in Chloroflexi. In this study, we analyze 16S rRNA gene sequence, quantitative PCR (qPCR), metagenomic and metatranscriptomic data that identify metabolic features of the Chloroflexi associated with their selection and subsistence over million-year timescales in oxic and anoxic deep-sea clay at multiple deep-sea sampling locations. The data reveal differences in metabolism between aerobic and anaerobic Chloroflexi persisting under oxic and anoxic energy-limited conditions for millions of years below the abyssal seafloor.

## Results

### Sedimentation rate and pore water geochemistry

We analyzed long (ca. 30 m) sediment cores from three deep-sea coring locations sites from R/V Knorr expedition KN223 in the Western North Atlantic, which were comprised of abyssal clay (Fig. 1A). Dissolved oxygen is present throughout the clay at sites 11 and 12, with concentrations at the water-sediment interface just below that of the overlying water (approximately 300 μM) and gradually decreasing with sediment depth (Fig. S1). Drawdown of O_2_ with sediment depth at these sites reflects oxidation of organic matter by aerobic microorganisms (Vuillemin et al., 2019). In contrast, dissolved oxygen at site 15 is restricted to the top mm of sediment. Sedimentation rate at both oxic sites sampled is estimated to be 1 m per million years (D’Hondt et al., 2015). Thus, the clay sampled at the deepest portion of the cores analyzed from both oxic sites (site 11, site 12) is about 30 million years old (Vuillemin et al., 2019). In contrast, anoxic site 15 is characterized by a mean sedimentation rate of about 3 m per million years (Fig. 1A). Given this rate, the deepest sediment sampled from the anoxic site 15 at 29 meters below seafloor (mbsf) is between 8 and 9 million years old.

**Figure 1.**
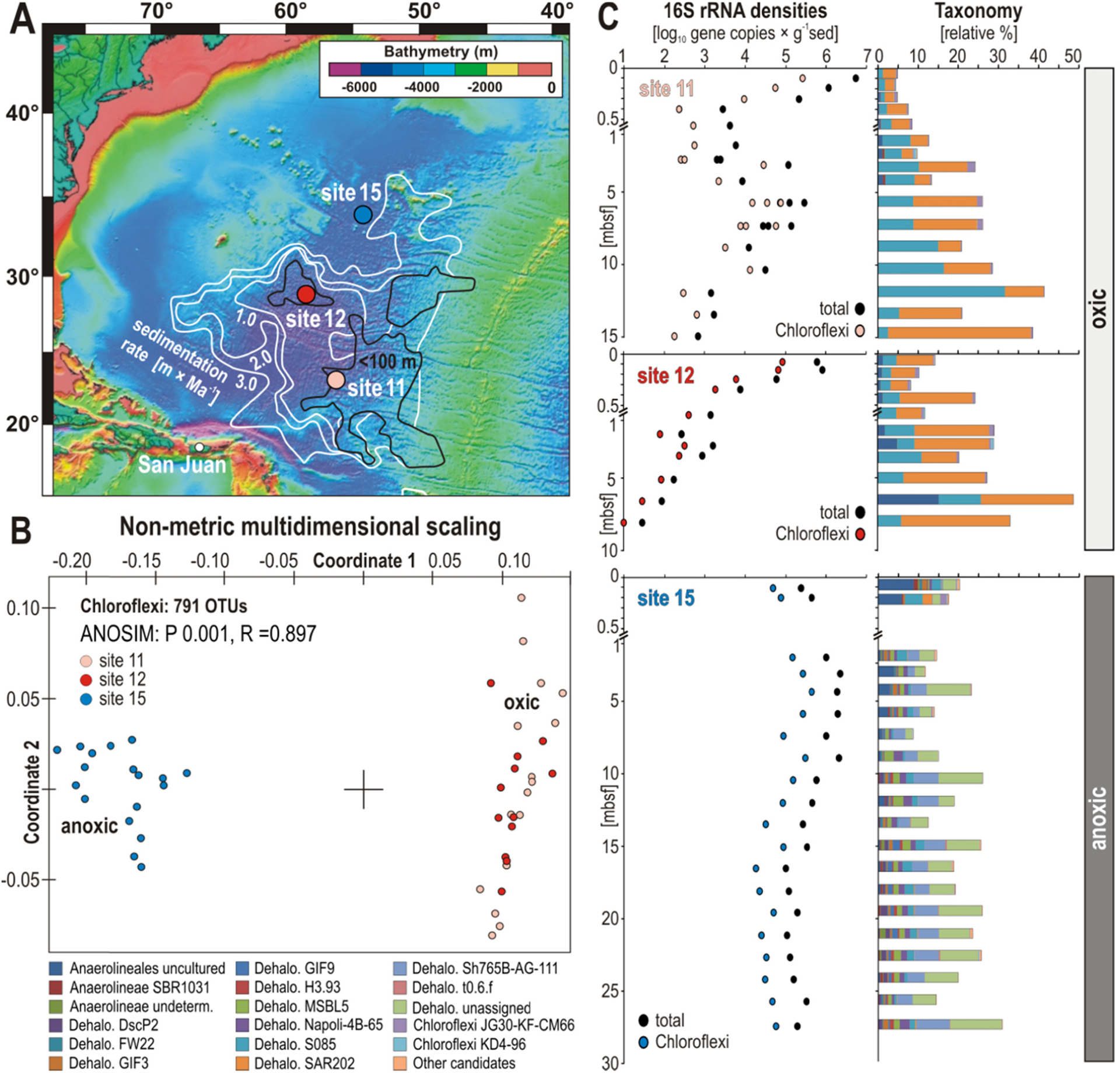
Map of sampling sites with bathymetry and sedimentation rate, and characterization of Chloroflexi in terms of beta diversity, density and taxonomy at the three sites. (**A**) Map of subseafloor sedimentation rate (after D’Hondt et al., 2015) with location of the three sampling sites. (**B**) Non-metrical dimensional scaling (NMDS) plot based on all OTUs assigned to Chloroflexi across the three sites. (**C**) Quantitative polymerase chain reaction (qPCR) of total (black dots) and qPCR-normalized to Chloroflexi (colored dots) 16S rRNA genes [gene copies × g^−1^ sed], and Chloroflexi taxonomic assemblages [relative %].

### Diversity, density and taxonomy of 16S rRNA genes

Four biological replicates of 16S rRNA genes (e.g. from separate DNA extractions) were sequenced at each sediment horizon from the anoxic sediment at site 15, which was compared to 16S rRNA genes from the two separate coring locations (site 11 and 12) that contain oxic clay. Based on 3711 OTUs obtained across all samples, non-metric multidimensional scaling (NMDS) analysis groups samples from the oxic sites separately from those of the anoxic site (Fig. S2) showing that oxygen conditions in the sediment had a statistically significant effect on community structure (ANOSIM: P 0.001, R = 0.897). All three sites exhibit similar values of total OTU richness (approximately 1500 at each site), Chloroflexi being the richest phylum (Fig. S2). Taxonomic assignment of 16S rRNA gene amplicons resulted in 791 Chloroflexi OTUs. Of these 791 OTUs, 555 are assigned to the class Dehalococcoidia, 136 are assigned to Anaerolineae, and the remaining 100 OTUs are in 9 different classes (Figs. S3-S4). Based on NMDS analysis of these 791 OTUs, the distribution of Chloroflexi between oxic and anoxic sediment was found to be statistically significant (ANOSIM: P 0.001, R = 0.897) (Fig. 1B).

At both oxic sites, the 16S rRNA gene densities (in copies × g^−1^ wet sediment) are highest in the shallow subsurface, respectively 10^7^ and 10^6^ copies × g^−1^ wet sediment, and decrease by 3 orders of magnitude within the underlying 0.5 mbsf (Fig. 1C). At oxic site 12, our limit of detection (10^2^ × 16S rRNA gene copies per DNA extraction) is reached at ∼8 mbsf, whereas at oxic site 11, the density of 16S rRNA genes increases by two orders of magnitude at ∼6 mbsf before reaching the detection limit at 15 mbsf. Similar to the procedure described by Lloyd et al. (2020), we normalized the fractional abundance of the Chloroflexi 16S rRNA abundance to quantitative values using the qPCR determined 16S rRNA gene concentrations as described previously (Vuillemin et al., 2019). The 16S rRNA gene density of Chloroflexi at both oxic sites sampled decreases constantly with depth, reaching the limit of detection at 15 mbsf at site 11 and at 10 mbsf at site 12. In contrast, the relative abundance of 16S rRNA genes assigned to Chloroflexi increases gradually at both oxic sites from <10% at the surface to about 40% in the deepest samples (Fig. 1C). The corresponding taxonomic assemblage is mainly composed of candidate clade S085 and SAR202 among Dehalococcoidia and uncultured Anaerolineales (Fig. 1C).

As implied by the NMDS analysis (Fig. 1B), the community composition at the anoxic site was significantly different by comparison. In contrast to the declining abundance of subseafloor Chloroflexi at oxic sites 11 and 12, at the anoxic site 15 the 16S rRNA gene density of Chloroflexi increases from 10^5^ copies at the seafloor surface up to two orders of magnitude in the deeper subsurface samples down to 2 mbsf, spanning ca. 1 million years of depositional history (Fig. 1C). Below this depth, the 16S rRNA gene density of Chloroflexi decreases by an order of magnitude and remains constant down to the deepest sampled interval at 29 mbsf. Throughout the anoxic sediment column, Chloroflexi account for about 20% of the complete taxonomic assemblage (Fig. 1C). Within the uppermost 0.3 mbsf at the anoxic site 15, Chloroflexi communities are similar to those of the oxic sites, namely candidate clades S085 and SAR202 and diverse uncultured Anaerolineales. However, below this depth at the anoxic site 15, the assemblages shift to presumed anaerobic candidates (e.g. GIF9, MSBL5, Sh7765B-AG-111) and uncultivated clades related to *Dehalococcoides* (Figs. 1C, S4). We acknowledge that primers targeting the V4-V6 regions as well as the average number of 16S rRNA gene copies in genomes of Chloroflexi (2.2 ± 1.2 copies) can potentially result in quantitative biases between the clades mentioned (Větrovský and Badrian, 2013).

### Metabolic potential in metagenomes

As described in several previous studies (Orsi et al., 2018; Ortega-Arbulù et al., 2019; Vuillemin et al., 2019; Orsi et al., 2020a), we used a bioinformatics pipeline involving a large aggregated genome database of predicted proteins including subsurface MAGs and SAGs (see Materials and Methods) for annotating putative functions of open reading frames (ORFs) in metagenomes and metatranscriptomes from particular “higher-level” (e.g. phyla or class level) taxonomic groups of microorganisms (e.g. Chloroflexi). This approach does not allow us to draw conclusions about specific populations within those groups (e.g. “MAGs” and “SAGs”), but does allow us to draw conclusions about metabolic traits derived specifically from higher-level taxonomic groups (Orsi et al., 2020a), such as Chloroflexi-related microorganisms given the ORF annotations provided. As is the case in all metagenomic studies, the incomplete nature of genomes in databases, together with the lower representation of sequenced genomes from candidate clades compared to cultured ones, make it likely that our pipeline misses annotation of ORFs that are derived from as-of-yet unsequenced Chloroflexi genomes that are not yet in databases. We acknowledge that some genes in databases annotated as being present in Chloroflexi MAGs might not actually derive from Chloroflexi chromosomes, but have been assigned to bins according to criteria that differ from study to study.

To assess the diversity of Chloroflexi recovered in metagenomes from the oxic and anoxic sites, we performed a phylogenetic analysis of the RNA polymerase sigma factor (*RpoD*) gene (Fig. 2A). We chose this gene for three reasons: (1) It is a universal single copy gene (Kawai et al., 2014); (2) the *RpoD* gene has a faster evolutionary rate compared to 16S rRNA genes and can be used to distinguish taxa at a relatively finer taxonomic level in phylogenetic analyses (Ghyselinck et al., 2013; Gupta et al., 2013); and (3) the *RpoD* gene was detected in Chloroflexi annotated contigs from metagenomes created at both oxic and anoxic sites, allowing for a comparative phylogenetic analysis of Chloroflexi in the metagenomes across sites. Phylogenetic analysis of the *RpoD* gene revealed the presence of several Chloroflexi clades among the SAR202 cluster in the oxic site metagenomes, and anaerobic clades of Dehalococcoidia in the anoxic site metagenomes (Fig. 2A), with affiliation to the recently published Chloroflexi genomes from deep-sea sediments (Fincker et al., 2020). Compared to the highly diverse candidate clades identified in the 16S rRNA gene taxonomy (Fig. S4), the phylogenetic analysis of *RpoD* gene indicated that the SAR202 clade and anaerobic deep-sea Dehalococcoidia were dominating the metagenomes at the oxic and anoxic sites, respectively. *RpoD* encoding genes from Chloroflexi were only detected in the metagenomes, no *RpoD* encoding transcripts from Chloroflexi were detected in the metatranscriptomes.

**Figure 2.**
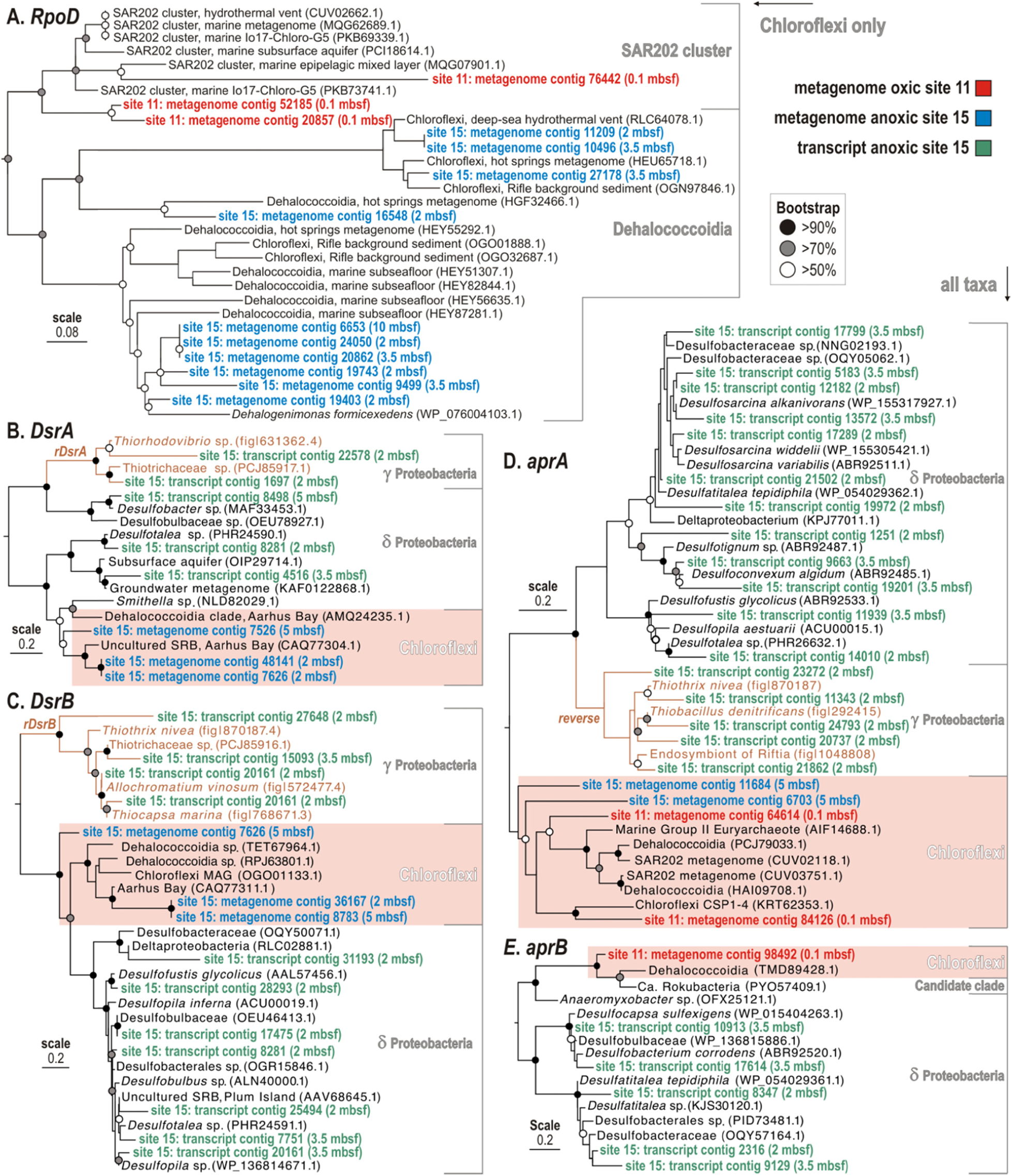
Phylogenetic analysis of predicted proteins encoded by selected marker genes in the metagenomes and metatranscriptomes from oxic and anoxic sites, based on RAxML using BLOSUM62 as the evolutionary model. **(A)** Phylogeny of Chloroflexi RNA polymerase sigma factor (*RpoD*) ORFs (619 aligned amino acid sites). **(B)** Phylogeny of all *DsrA* and *rDsrA* ORFs detected (466 aligned amino acid sites). **(C)** Phylogeny of all *DsrB* and *rDsrB* ORFs detected (433 aligned amino acid sites). Panel**s (D)** and **(E)** Phylogenies of all *aprA* and *aprB* ORFs detected (754 and 157 aligned amino acid sites, respectively). All phylogenies were conducted with 100 bootstrap replicates, circles on the nodes indicate support values (black > 90%, gray > 70%, white >50%).

For both oxic sites, the number of unique ORFs in metagenomes assigned to Chloroflexi is highest in surface sediment and decreases by exponentially with increasing sediment depth (Fig. 3A). This mirrors the trend of declining Chloroflexi abundance with depth at the oxic sites based on qPCR and 16S rRNA gene sequencing (Fig. 1C). Because the multiple displacement amplification (MDA) step precludes a discussion of relative abundance of genes, we focus on presence or absence of unique ORFs assigned to Chloroflexi. In the shallow oxic sediment, Chloroflexi ORFs could be assigned to clusters of orthologous genes (COGs) involved in energy conversion, transport and metabolism of amino acids, carbohydrates, and coenzymes (Fig. 3A). As the detection of annotated protein-encoding ORFs decreases with depth, COG categories only include nucleotide, coenzyme and lipid transports and metabolisms. The relative detection of metabolic pathways and functional gene categories assigned to Chloroflexi indicate mainly aerobic respiration via cytochrome C oxidase and restricted potential for the tricarboxylic acid (TCA) cycle. Energy production relates to ATP synthase, NADH-quinone and -flavin reductase. Dehalogenases, peptidases, lipases, glycosidases, and oxidation of volatile fatty acids (VFAs) were all detected.

**Figure 3.**
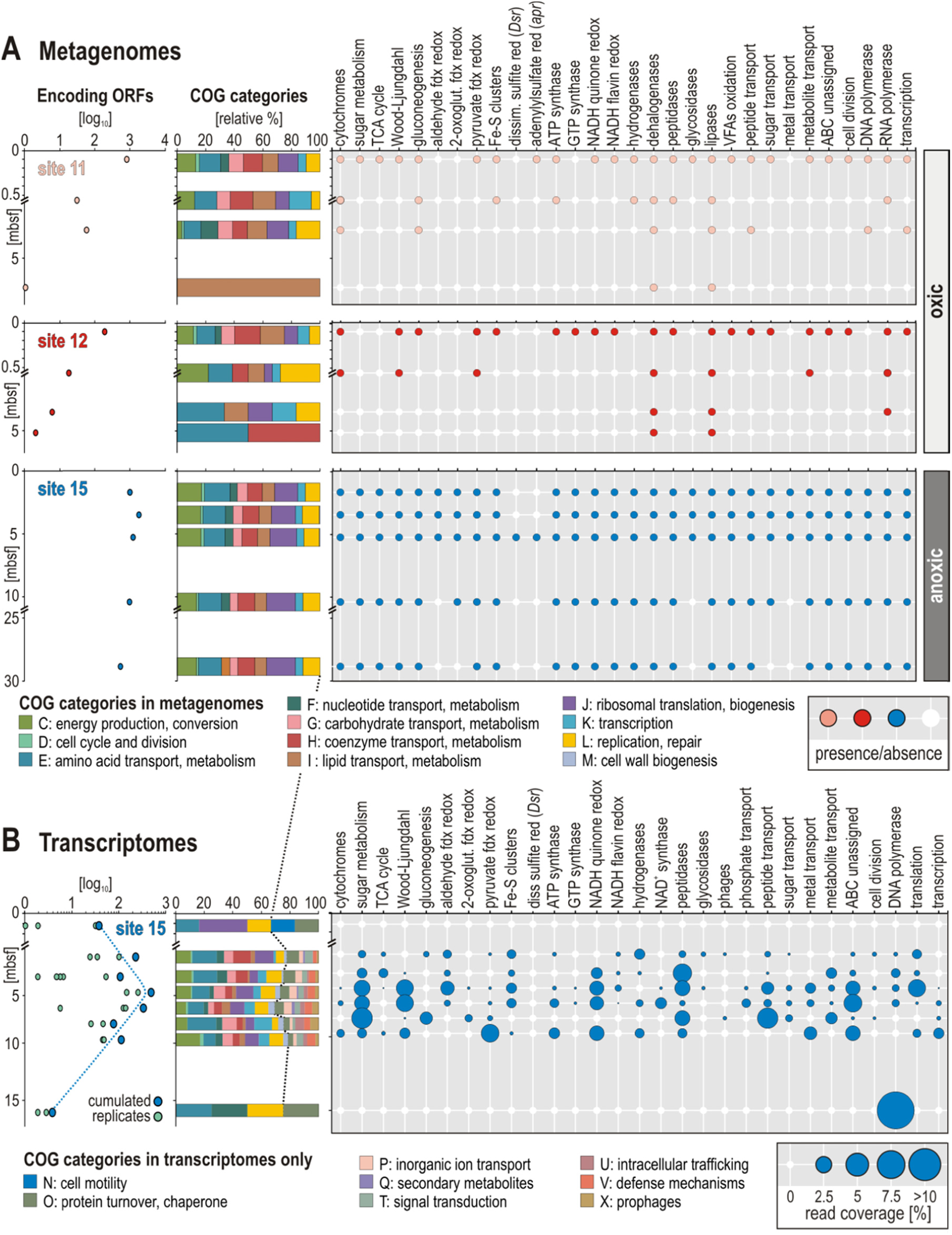
Open reading frames (ORFs) assigned to Chloroflexi, their COG categories and metabolic potential obtained from metagenomes and metatranscriptomes. (**A**) Number of annotated protein-encoding ORFs obtained from metagenomes and attr**i**buted to Chloroflexi with sediment depth; relative abundances of different COG categories and metabolic functions obtained from metagenomes at the three sites. (**B**) Number of annotated protein-encoding ORFs obtained from the metatranscriptomes and attributed to Chloroflexi with sediment depth; relative abundances of different COG categories and metabolic functions that are expressed at the anoxic site.

At the anoxic site, detection of annotated protein-encoding ORFs assigned to Chloroflexi in metagenomes remains constants with depth throughout the entire 29 m core sediment sequence (Fig. 3A). The main corresponding COG categories are energy production, transport and metabolism of amino acids, carbohydrates, coenzymes and lipids, but also include cell biogenesis, transcription and replication. The metabolic pathways from which the ORFs assigned to Chloroflexi are involved are related to sugar metabolism, the TCA cycle and W-L pathway. ORFs annotated as energy metabolism (electron carrier) genes from the Chloroflexi were identified at the anoxic site, including pyruvate and aldehyde ferredoxin oxidoreductase, NADH-quinone and -flavin oxidoreductase.

Also related to energy production, six Chloroflexi ORFs annotated as the alpha and beta subunits of the *apr* and *Dsr* genes were detected between 2 and 5 mbsf in the metagenomes at the anoxic site 15 (Figs. 2B-2E), which are affiliated to Chloroflexi known from published SAG (Wasmund et al., 2014). No *aprAB* or *DsrAB* encoding transcripts from Chloroflexi were detected in the metatranscriptomes. Instead, numerous *aprAB* or *DsrAB* encoding transcripts were detected at anoxic site 15, mostly from known groups of sulfate-reducing Deltaproteobacteria (Müller et al., 2015). At the oxic site 11, at a depth of 0.1 mbsf, *aprA* and *aprB* subunits were found to be encoded in metagenomes with affiliation to Chloroflexi ORFs (Figs. 2D-2E), but again no transcripts were detected with expression of these genes. Thus, the metabolic potential for the associated sulfur cycling in Chloroflexi is present at all sites, but its transcription was below detection in all cases. This confirmed metabolic potential for dissimilatory sulfate reduction among clades of Dehalococcoidia, whereas the absence of the related ORFs in our metatranscriptomes suggested that these genes are not actively transcripted by Chloroflexi in the anoxic sediment.

### Metabolic potential in metatranscriptomes

At the anoxic site, the number of unique annotated protein-encoding ORFs assigned to Chloroflexi increases by two orders of magnitude from the seafloor surface to 5 mbsf. Below this depth, the number of unique ORFs assigned to Chloroflexi steadily decreases with depth until 15.9 mbsf. Below this depth, the only ORFs annotated were those assigned to groups of known contaminants from molecular kits including those from human skin and soil (Salter et al., 2014). Many of these same groups are common laboratory contaminants found in dust samples from our lab in 16S rRNA gene surveys (Pichler et al., 2018), and include *Pseudomonas, Rhizobium, Acinetobacter*, and *Staphylococcus*. We interpreted the sudden dominance of ORFs with similarity these common contaminants below 15.9 mbsf to be indicative of our limit of RNA detection, whereby a lower amount of extracted RNA from the *in situ* active community becomes overprinted by background “noise” from contaminating DNA.

In metatranscriptomes from 0.1 mbsf at the anoxic site 15, COG annotations of ORFs assigned to Chloroflexi only included biogenesis, DNA replication, protein turnover, and motility (Fig. 3B). Interestingly below 0.1 mbsf, there was no longer any gene expression of ORFs encoding proteins involved in motility. This indicates that motility of Chloroflexi was possibly restricted to the upper sediment layers, with starvation-induced loss of this function below (Wei and Bauer, 1998; Zhu and Gao, 2020). Several COG categories increased in relative abundance in the metatranscriptomes of Chloroflexi at deeper depths, which included ORFs involved in cellular defense, prophages, motility and secondary metabolites (Fig. 3B). These COG categories were expressed in the metatranscriptomes but were not detected in the metagenomes, indicating a relatively high level of expression. Rather, the increasing relative abundance of these COG categories in the metatranscriptomes may correspond to activities that increase in relative abundance over time since burial. While a bias by MDA amplified metagenomes cannot be completely ruled out, it is unlikely that MDA would be biased against entire COG categories. This is consistent with a higher number of unique annotated ORFs assigned to Chloroflexi deeper in the subsurface, which is an indication of their increased activity in the older sediments (Fig. 3B). In the metatranscriptomes, Chloroflexi ORFs encoding proteins involved in metabolic pathways could be assigned to sugar metabolism, and the W-L pathway. The deepest ORFs assigned to Chloroflexi at 15.9 mbsf are involved in cellular maintenance activities, such as DNA repair and protein turnover. Expression of Chloroflexi ORFs involved in energy production annotated as aldehyde and pyruvate ferredoxin oxidoreductase, NADH-quinone oxidoreductase (Nuo), ATP synthase and NADH-flavin oxidoreductase were also detected (Fig. 3B).

An NMDS analysis based on all annotated protein-encoding ORFs obtained from metagenomes and metatranscriptomes (ANOSIM: P 0.001, R = 0.56), as well as those exclusively assigned to Chloroflexi (ANOSIM: P 0.005, R = 0.34), clearly separates all of the metagenome and metatranscriptome samples from both oxic sites from those recovered from the anoxic site (Fig. 4). This demonstrates that the metabolic potential is significantly different, not only within the entire microbial community, but between aerobic and anaerobic Chloroflexi as well. Because the sediment samples are millions of years old, selection for specific metabolisms has occurred in these different Chloroflexi over exceptionally long timescales.

**Figure 4.**
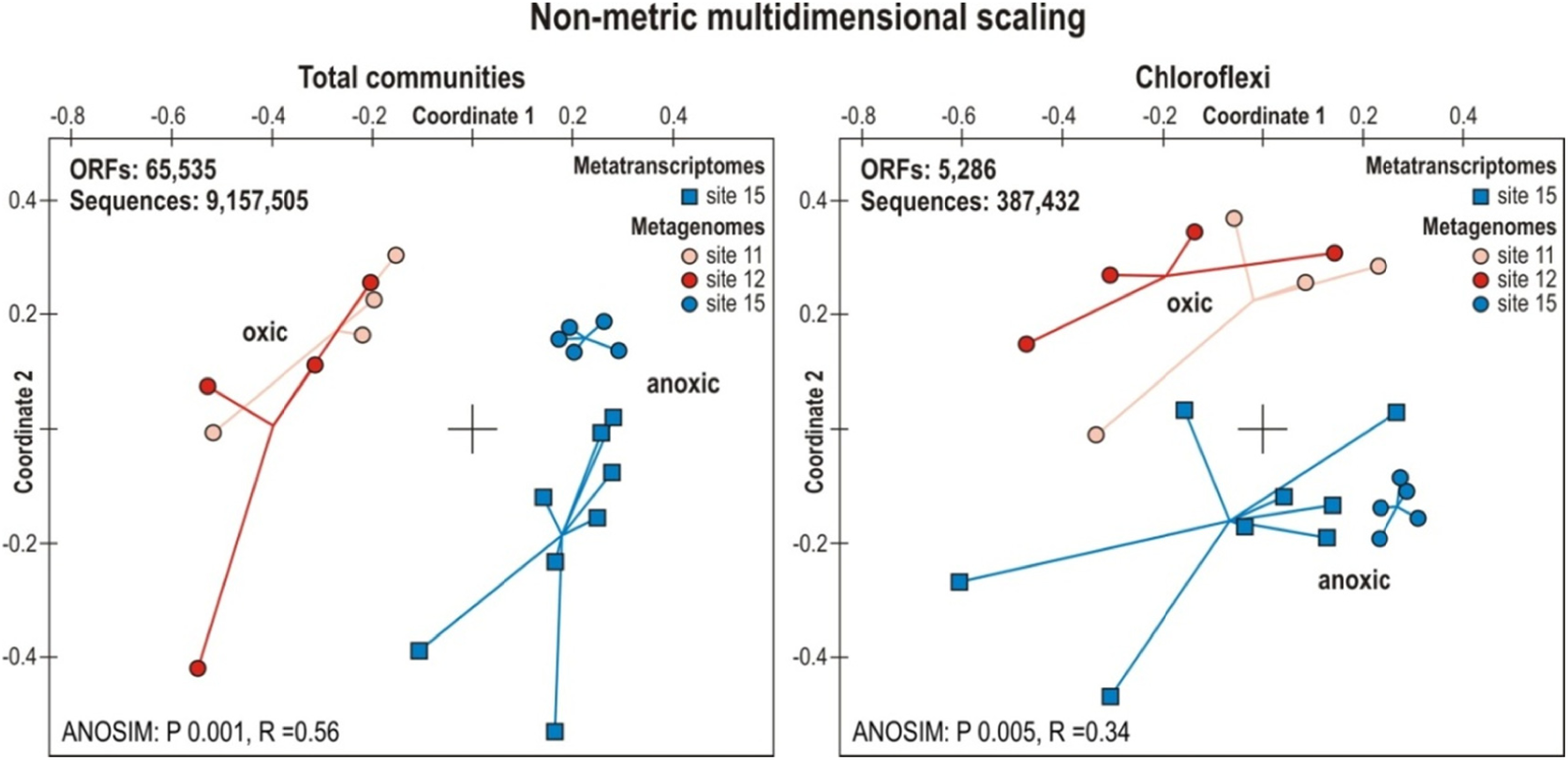
Non-metrical dimensional scaling (NMDS) plot for all three sites based on metagenomes and metatranscriptomes. (**Left**) NMDS plot for all three sites based on all ORFs from the metagenomes (circles) and metatranscriptomes (squares) for the entire microbial communities. (**Right**) NMDS plot for all three sites based on annotated protein-encoding ORFs from the metagenomes (circles) and metatranscriptomes (squares) assigned to Chloroflexi.

From the oxic site samples, no metatranscriptomes of sufficient quality could be obtained. We attempted to extract RNA and make metatranscriptome libraries from the samples, but in every sample the only recovered ORFs annotated were those assigned to groups of known contaminants from molecular kits (Salter et al., 2014) and dust samples from our lab (Pichler et al., 2018). We interpreted the dominance of ORFs with similarity to these common contaminants to be a clear indication of contamination, since the metagenomes and 16S rRNA gene datasets showed a completely different community structure that was dominated by the Chloroflexi. In none of the metatranscriptomes from the oxic site could we retrieve any ORFs annotated to Chloroflexi genomes, indicating that the gene expression and overall activity of Chloroflexi at the oxic sites were very low.

Phylogenetic affiliation of ORFs from the metatranscriptomes encoding the *DsrAB* genes (Fig. 2B-C) pointed to Deltaproteobacteria as the main active sulfate-reducing bacteria subsisting in the anoxic subseafloor clay at site 15, whereas a smaller number of expressed ORFs were affiliated with Gammaproteobacteria reverse *DsrAB* (*rDsr*) involved in sulfide oxidation (Loy et al., 2009). Phylogenetic analysis of the ORFs encoding *aprA* and *aprB* proteins identified in our metagenomes and metatranscriptomes pointed to Delta- and Gammaproteobacteria as the main clades actively expressing these genes in the anoxic subseafloor at site 15 (Figs. 2C-2D). Five ORFs assigned to *aprAB* genes in metagenomes from oxic site 11 and anoxic site 15 were affiliated to Chloroflexi MAGs among the SAR202 cluster (Mehrshad et al., 2018), revealing metabolic potential for *aprA* and *aprB* gene in both oxic and anoxic subseafloor sediment. However, same as for the *DsrAB* genes encoded by the subseafloor Chloroflexi, there was no detection for active transcription of these genes by Chloroflexi in our metatranscriptomes.

## Discussion

We present new data from million year old sediments that provide new insights into how life survives under energy-limited conditions sediments, both in oxic and anoxic sediments, over million-year timescales. Subseafloor life in oxic sediments is energetically limited by the availability of electron donors (organic matter), whereas in anoxic sediments it is energetically limited by the availability of high energy terminal electron acceptors. Since Chloroflexi constitute a substantial part of the subseafloor communities under both settings, we investigated metabolic potentials and strategies that may allow them to actively survive under extreme energy limitation and persist in subseafloor sediment for millions of years after burial.

### Chloroflexi in oxic deep-sea clay

At both oxic sites, the Chloroflexi were dominated by the SAR202 clade, and the 16S rRNA gene density of Chloroflexi decreased exponentially with depth (Fig. 1C), indicating net death over this interval of burial. The clade SAR202 dominates in abyssal oxic sediment communities in many locations, indicating it is relatively well-suited to cope with the otherwise unfavorable (e.g. energy-limited) conditions for life (Durbin and Teske, 2011; Mehrshad et al., 2018). Phylogenetic analysis of the Chloroflexi *RpoD* gene proteins from confirmed taxonomic affiliation of protein-encoding ORFs to the SAR202 clade in metagenomes from the oxic sites (Fig. 2A). In metagenomes from the oxic sites, we identified metabolic potential for aerobic respiration via cytochrome C oxidase (*COX*) (Fig. 3A). As previously described for the SAR202 clade, evidence for metabolic use of refractory organic matter (Landry et al., 2017) includes detection of genes that code for haloalkane and haloacetate dehalogenases, peptidases and oxidation of VFAs via acyl-CoA dehydrogenase – all of which were detected from the Chloroflexi at the oxic site metagenomes, indicating their potential to survive off of consuming refractory organic matter in the million year old clay (Fig. 3A). The fact that the extractable RNA for both oxic sites was below detection indicates that the aerobic SAR202 Chloroflexi dominating at the oxic sites have a lower level of activity, compared to the anaerobic Chloroflexi that were subsisting at the anoxic site that had detectable levels of transcriptional activity that actually increase with depth (Figs. 3A-3B). According to Mehrshad et al. (2018), some SAR202 Chloroflexi may use the *apr* gene during aerobic sulfate metabolism, possibly in the oxidation of sulfite or thiosulfate produced during the breakdown of organic sulfur molecules. The SAR202 in the oxic clays also contain this *apr* gene (Fig. 2D), which is related to *apr* genes from some Euryarchaeota (Mehrshad et al. 2018), but no metatranscriptomes from the oxic clays could be produced presumably due to the ultra-low metabolic activities of the cells. Thus, it remains unknown whether the subseafloor SAR202 Chloroflexi expressed this *apr* gene or not, and whether they are engaged in sulfite or thiosulfate oxidation. Nevertheless, the presence of this gene in the metagenomes indicates the metabolic potential for such processes.

### Chloroflexi in anoxic deep-sea clay

The increase in gene density below the seafloor (Fig. 1C) and number of unique expressed ORFs (Fig. 3) suggest that Chloroflexi proliferate in the deep anoxic clay and experience net growth over the top few meters of sediment. The Chloroflexi 16S rRNA genes at the seafloor surface at anoxic site 15 were predominantly affiliated with uncultivated Anaerolineales and the Dehalococcoidia. Their most closely related cultivates (Fig. S4), i.e. *Anaerolinea, Leptolinea* (Yamada et al., 2006), *Longilinea* (Yamada et al., 2017), *Pelolinea* (Imachi et al., 2014), and *Flexilinea* (Sun et al., 2016), are known to mainly ferment sugars but, unlike their main class representative *Dehalococcoides mccartyi* (Löffler et al., 2013), they do not grow on VFAs or alcohols as electron donors or haloorganic electron acceptors. The taxonomic composition of the Chloroflexi assemblage transitioned away from the SAR202 at the surface of site 15 towards anaerobic clades of Dehalococcoides with increasing depth (Fig. 1C). Phylogenetic analysis of the Chloroflexi *RpoD* encoded proteins confirmed the presence of Dehaloccoidia clades in the metagenomes from the anoxic site (Fig. 2A). These metagenomes further revealed the presence of Chloroflexi assigned ORFs annotated to proteins involved in sugar metabolism, the W-L pathway, pyruvate ferredoxin oxidoreductase, ATP synthase, and NADH-quinone oxidoreductase (Fig. 3A). The NMDS plots for 16S rRNA genes (Fig. 1B) and metagenomes (Fig. 4) significantly separate oxic and anoxic samples, indicating that the presence or absence of O_2_ exerts very strong selection pressure (ANOSIM: P 0.001, R = 0.897; P 0.001, R = 0.56).

The observation that metatranscriptomes at the anoxic site recovered relatively high numbers of ORFs annotated to Chloroflexi, whereas no ORFs from metatranscriptomes could be annotated to Chloroflexi from the oxic site samples, indicates that anaerobic Chloroflexi in the anoxic subseafloor clay had a generally higher level of activity compared to aerobic Chloroflexi in the oxic subseafloor clays (Figs. 2A-2B). At the anoxic site at a depth of 0.1 mbsf, Chloroflexi ORFs in metatranscriptomes were involved in translation, replication and protein turnover (Fig. 3B). As the density of 16S rRNA genes (Fig. 1C) and number of annotated protein-encoding ORFs (Fig. 2B) increases with sediment depth, the number of different COG categories in the metatranscriptomes assigned to Chloroflexi increases, suggesting that the metabolic activity of the anaerobic Chloroflexi increases with sediment depth at the anoxic site. These additional COG categories from the metatranscriptomes included transcription, energy production, cellular defense, prophages, and production of secondary metabolites. In the deepest metatranscriptome at 15.9 mbsf, the only annotated protein-encoding ORFs that were assigned to Chloroflexi were involved in DNA repair, nucleotide transport, metabolism and protein turnover, which are indicative of cellular maintenance processes in sediments that have an estimated age of 5 million years (Fig. 3B).

The metatranscriptomes from the anoxic site point to fermentation and acetogenesis as key metabolic mechanisms for anaerobic subseafloor Chloroflexi, which was already predicted from genome analysis of subseafloor Chloroflexi (Sewell et al., 2017; Fincker et al., 2020). The detection of ORFs assigned to Chloroflexi with similarity to the methyl-viologen Fe-S reducing and F420 non-reducing Ni-Fe hydrogenases (*MVHs*) indicated the potential for production of molecular hydrogen (Vignais et al., 2001; Shafaat et al., 2013) (Fig. 5). The utilization of a partial W-L pathway in anaerobic carbon metabolism has been shown in the cultivated Chloroflexi strain *Dehalococcoides mccartyi*, which uses a partial W-L pathway for C1 compound assimilation from either CO_2_ reduction or acetate oxidation (Zhuang et al., 2014; Fincker et al., 2020). Expression of Chloroflexi ORFs annotated as pyruvate ferredoxin oxidoreductase (Fig. 3B) may provide a link between the W-L pathway and other anabolic pathways (Wasmund et al., 2014). Here, the ORFs assigned to Chloroflexi with similarity to W-L pathway proteins that are actively expressed included formate dehydrogenase (*fdh*), methylenetetrahydrofolate reductase (*MTHFR*), carbon monoxide dehydrogenase (*CODH*) and acetyl-CoA synthase (*cdhA*) (Fig. 5). Many homoacetogenic bacteria use the Rnf complex to generate a chemiosmotic gradient of Na^+^ ions in order to synthesize ATP at the membrane (Peters et al., 2016; Schuchmann and Müller, 2019). However, Dehalococcoidia appear to use the NADH-quinone oxidoreductase (Nuo) to create a proton gradient in order to generate a chemiosmotic potential for ATP synthesis (Wasmund et al., 2017). Nuo yields reduced energy compared to the flavin-dependent oxidoreductase (Buckel and Thauer, 2018). ORFs with similarity to Nuo and ATP synthases are expressed by the Chloroflexi in greater abundances down to 10 mbsf (Fig. 3B), which potentially related to active ATP production via chemiosmotic proton pumping with Nuo. Ferredoxin and flavodoxin (Fig. 5) are expressed by the Chloroflexi at these same depths and are likely involved as electron carriers in cellular redox reactions (Peters et al., 2016; Buckel and Thauer, 2018).

**Figure 5.**
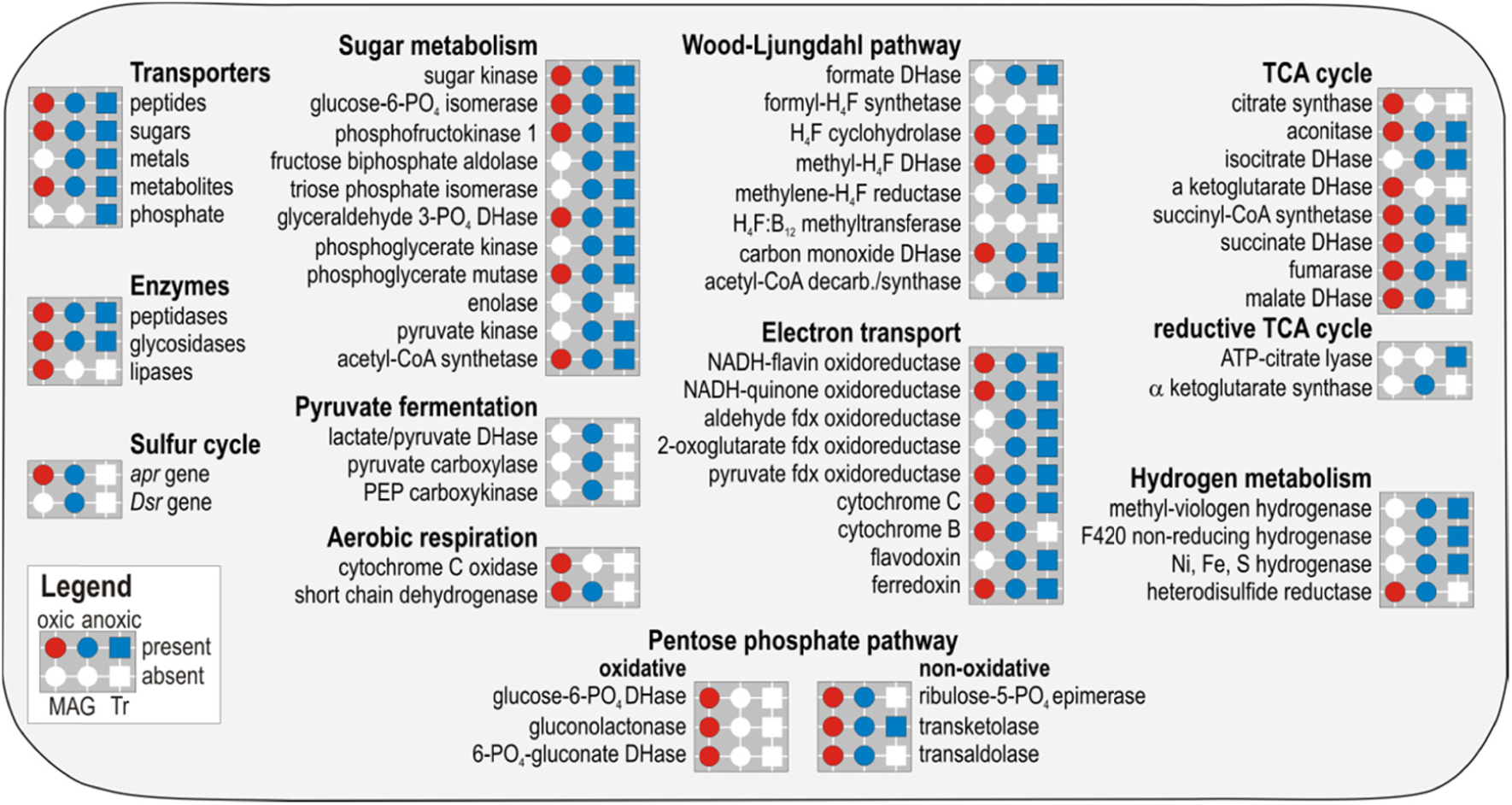
Presence and absence of Chloroflexi assigned predicted proteins in metagenomes and metatranscriptomes. Presence (colored) or absence (white) of metabolic functions (circles) and expressed genes (squares) identified in the metagenomes and metatranscriptomes at the oxic (red) and anoxic (blue) sites across all depths sampled. Presence/absence of substrate transporters, catabolytic enzymes, *apr* and *Dsr* genes are shown on the left hand side. Metabolic pathways listed correspond to sugar metabolism, pyruvate fermentation, aerobic respiration, Wood-Ljungdahl pathway, electron transport, tricarboxylic acid cycle, hydrogen metabolism, and oxidative and non-oxidative pentose phosphate pathway. The complete list of genes with coverage is available as Tables S2-S3.

Additional Chloroflexi ORFs with similarity to W-L pathway proteins were also present in the metagenomes, which were annotated as methylene-tetrahydrofolate dehydrogenase/cyclohydrolase (*DHCH*), and acyl-CoA dehydrogenase (*ACAD*) (Fig. 5). This indicates that some of the anaerobic Chloroflexi in the anoxic subseafloor clay have metabolic potential for β-oxidation of VFAs. Chloroflexi ORFs annotated as acetate kinase, which is the enzyme that yields one ATP during substrate level phosphorylation (SLP) in (homo)acetogenic bacteria (Schuchmann and Müller, 2019), was detected in the metagenomes (Table S2) but not in the metatranscriptomes (Table S3). This might indicate ATP production via chemiosmotic ion gradients at the membrane as a preferential source of energy production (as opposed to dephosphorylation of acetyl-phosphate) by anaerobic Chloroflexi in these million year old anoxic clays.

### Survival mechanisms of Chloroflexi in oxic and anoxic subseafloor clay

The organic matter present at very low concentrations in abyssal clay is dominated by amide and carboxylic carbon and generally proteinaceous (Estes et al., 2019). We infer that Chloroflexi actively utilize this organic matter in the anoxic subseafloor clays, based on the relatively high expression levels of peptide transporters and peptidases (Fig. 3B). During fermentation of amino acids, hydrogen is produced (Buckel and Barker, 1974; Barker, 1981), which can ultimately be cycled inside the cell and channeled into the partial W-L pathway (Wiechmann et al., 2020). Amino acid substrates are possibly the most important of all fermentation substrates in the subseafloor because cells subsist in these environments on necromass (Bradley et al., 2019), and most of cellular material is protein by weight (Orsi et al., 2020b). Thus, the proteinaceous components of deep-sea clay (Estes et al., 2019) likely serve an important role in sustaining the activity of anaerobic Chloroflexi seen here, possibly via the fermentation of amino acids, for millions of years after they are buried below the seafloor.

Such intracellular H_2_ recycling by homoacetogenic Chloroflexi in the subseafloor has been shown recently in a genomic analysis (Fincker et al., 2020), whereby a higher energy efficiency of this metabolism confers a fitness advantage for subseafloor homoacetogens including Chloroflexi. If true, this may imply that such acetogens do not require an H_2_ producing syntrophic partner (Wiechmann et al., 2020). Rather, they make the H_2_ themselves via fermentation and then consume it via the W-L pathway, which likely helps to recycle NADH and ferredoxin pools, leading to a higher ATP yield via chemiosmotic pumping of ion gradients at the membrane (Schuchmann and Müller, 2019). The close affiliation of a clade containing many Chloroflexi assigned *rpoD* sequences from the metagenomes of anoxic site 15 with *RpoD* sequences from deep-sea Chloroflexi MAGs (Fig. 2A) that have such a homoacetogenic metabolism (Fincker et al., 2020), further supports the conclusion that many of the transcriptionally active Chloroflexi at anoxic site 15 are surviving via homoacetogenesis.

The genomic potential for dissimilatory sulfate reduction in uncultivated clades of Chloroflexi from marine sediments has been identified previously (Wasmund et al., 2017; Mehrshad et al., 2018), which could be an important metabolic feature of Chloroflexi to survive under the energy-limited conditions for long time periods below the seafloor. The detection of ORFs assigned to *aprAB* and *DsrAB* genes in the metagenomes from the anoxic site (Fig. 2B-2E), indicates that some of the Chloroflexi have potential for dissimilatory sulfate reduction in the anoxic subseafloor sediment. Dissimilatory sulfate reduction could be utilized by subseafloor Chloroflexi to promote increased protein synthesis and growth in the million year old sediments sampled here. However, expression of ORFs encoding the *aprA* and *DsrAB* proteins assigned to Chloroflexi were not detected in the metatranscriptomes from the same samples (Fig. 5), which suggests that dissimilatory sulfate reduction is not a major energy yielding activity under these conditions compared to the Deltaproteobacteria that, in contrast, expressed many ORFs of *DsrAB* and *aprAB* genes (Figs. 2B-2E). The most genes expressed by the anaerobic Chloroflexi at the anoxic site appear to be rather involved in a homoacetogenic lifestyle, which has been proposed to be H_2_-syntroph independent and involves intracellular re-oxidation of H_2_ for CO_2_ fixation via the W-L pathway (Finker et al., 2020). The gene expression data assigned to Chloroflexi from these million year old deep-sea clays indicate that this proposed metabolic mechanism is actively used by subseafloor anaerobic Chloroflexi, and is more expressed compared to Chloroflexi genes involved in dissimilatory sulfate reduction. Based on this, we speculate that the subseafloor Chloroflexi, that have a potential for dissimilatory sulfate reduction, avoid competing for sulfate with sulfate-reducing bacteria by applying a homoacetogenic lifestyle instead. Our data show that this is correlated with increased net growth over a ca. 1 million year interval in the upper 5 m of deep-sea clay sediment at anoxic site 15.

Our metatranscriptomes show that Deltaproteobacteria are the main sulfate-reducing bacteria that are presently active in the anoxic subseafloor (Figs. 2B-2E). ORFs expressing the *rDsrAB* and reversible *aprA* genes (Loy et al., 2009) further suggest that some Gammaproteobacteria are active in the samples (Fig. 2), but it remains unclear what *rDsr* expressing organisms (e.g. H_2_S oxidizing bacteria) would use as a terminal electron acceptor. However, since the *Dsr* and *rDsr* enzymes are potentially reversible (Loy et al., 2009; Müller et al., 2015; Orsi et al., 2016), the expression of the *rDsr* genes from the usually sulfide oxidizing Gammaproteobacteria may function in these sediments as *Dsr* genes and, in this case, the *Dsr* enzyme in organisms may actually be performing sulfate reduction instead of sulfide oxidation.

Metagenomes from the oxic sites 11 and 12 also have Chloroflexi assigned *aprAB* encoding genes, but those are derived from the SAR202 clade (Figs. 2D-2E). Similar to anoxic site 15, there was no detectable expression of *aprAB* encoding transcripts at the oxic sites. Moreover, there were no *DsrAB* genes assigned to Chloroflexi that were detected in metagenomes or metatranscriptomes from any of the oxic site samples (Figs. 2B-2C). Sequences of the *aprAB* genes from oxic subseafloor Chloroflexi SAR202 clade constitute a separate group (Figs. 2D-2E), which have been proposed to correspond to a sulfur-oxidizing version of the *aprA* genes (e.g. reversed reaction) instead of sulfate reduction, that some SAR202 clade organisms could use to oxidize sulfite derived from organosulfur compounds (Mehrshad et al., 2018). In addition, aerobic Chloroflexi from the SAR202 clade at the oxic sites mainly display metabolic potential for aerobic sugar metabolism and a partial TCA cycle, and the entire oxidative and non-oxidative pentose phosphate pathway (Fig. 5), implying a potential mechanism to store and consume sugars over millions of years. Chloroflexi assigned ORFs from the oxic site metagenomes annotated as carbon monoxide dehydrogenase, heterodisulfide reductase, and Nuo (Fig. 5) have been suggested previously to be signatures of an aerobic carboxidotrophy lifestyle (Carr et al., 2019; Cordero et al., 2019; Islam et al., 2019). It is possible that the aerobic Chloroflexi SAR2020 clade organisms dominating in the subseafloor oxic clay at the two sampled locations are using a similar aerobic carboxidotrophy metabolism. Oxidation of CO has been shown to be energetically favorable in cultivated bacteria (Cordero et al., 2019; Islam et al., 2019), which might be advantageous for the aerobic Chloroflexi subsisting in oxic subseafloor clay in the absence of abundant organic matter (Bradley et al., 2019).

## Conclusions

Our findings demonstrate that transcriptionally active anaerobic, homoacetogenic Chloroflexi proliferate in anoxic abyssal clay sediments that are 2 to 3 million-years old, and exhibit higher activity compared to aerobic Chloroflexi from oxic abyssal clays. The gene expression data support prior predictions of potential (homo)acetogenic lifestyle, independent of an H_2_-producing syntrophic partner, for some subseafloor Chloroflexi. A metabolic potential for aerobic oxidation of CO in the aerobic Chloroflexi might be related to their successful dominance in the oxic abyssal clay communities. Because Chloroflexi communities in the oxic subseafloor exhibit net death over the million year time scales (e.g. increasing depth below seafloor), we infer that the majority of clade SAR202 cells persist in this setting as dormant or non-growing metabolic states. In contrast, anaerobic Chloroflexi appear to maintain detectable levels of transcriptional activity in anoxic abyssal clay deposited up to three million years ago.

We propose that this difference in activity between oxic and anoxic sites is related to the nature of the energy limitation. The oxic abyssal clays are energetically limited by the availability of electron donors (organic matter), which is why there is ample availability of electron acceptors in the form of oxygen and nitrate, and also why oxic clays are red (because the iron is oxidized). In contrast, the anoxic abyssal clays have higher concentrations of electron donors in the form of organic matter and are energy-limited by electron acceptors – all of the higher energy oxygen and nitrate are consumed via respiration close to the sediment surface. Given the high energy nature of oxygen and nitrate as electron acceptors, one might expect aerobic Chloroflexi in oxic abyssal clays to have higher activities compared to anaerobic Chloroflexi in anoxic abyssal clays, but our findings show that this is not the case. Rather, it appears that the activity of homoacetogenic Chloroflexi in anoxic sediment is correlated with the availability of electron donors in the form of organic matter rather than the availability of higher energy electron acceptors like oxygen and nitrate. This hypothesis could be tested with additional sampling of deep-sea cores from other oxic and anoxic clays, for example in new deep drilling expeditions to the South Pacific and North Pacific Gyres.

## Materials and Methods

### Sampling expedition

All samples were taken during Expedition KN223 of the *R/V Knorr* in the North Atlantic, from 26 October to 3 December 2014. At site 11 (22°47.0′ N, 56°31.0′ W, water depth ∼5600 m), site 12 (29°40.6′ N, 58°19.7′ W, water depth ∼5400 m), and site 15 (33°29.0’ N, 54°10.0’ W, water depth 5515 m), successively longer sediment cores were retrieved using a multicorer (∼0.4 m), gravity corer (∼3 m) and the Woods Hole Oceanographic Institution (WHOI) piston-coring device (∼29 m). Additional details of sampling are published elsewhere (D’Hondt et al., 2015 and 2019). Dissolved oxygen concentrations in the core sections were measured with optical O_2_ sensors as described previously (D’Hondt et al., 2015). Sediment subcores were retrieved on the ship aseptically using end-cut sterile syringes and kept frozen at −80 °C without any RNA shield until extraction in spring 2018 in the home laboratory.

### DNA extraction, quantitative PCR, 16S rRNA genes

For site 11 and 12, total DNA was extracted from 10 g of sediment per sample as previously described (Vuillemin et al., 2019), whereas for site 15 we used 0.7 g of sediment per sample and diluted the final extracts 10 times in ultrapure PCR water (Roche). DNA templates were used in quantitative PCR (qPCR) amplifications with updated 16S rRNA gene primer pair 515F (5′-GTG YCA GCM GCC GCG GTA A -3′) with 806R (5′-GGA CTA CNV GGG TWT CTA AT -3′) to increase our coverage of Archaea and marine clades and run as previously described (Pichler et al., 2018). All qPCR reactions were set up in 20 µL volumes with 4 µL of DNA template and performed as previously described (Coskun et al., 2019). Reaction efficiency values in all qPCR assays were between 90% and 110% with *R*^*2*^ values >0.95% for the standards. For 16S rRNA gene library preparation, qPCR runs were performed with barcoded primer pair 515F and 806R. All 16S rRNA gene amplicons were purified from 1.5% agarose gels, normalized to 1 nM solutions and pooled. Library preparation was carried out according to the MiniSeq System Denature and Dilute Libraries Guide (Protocol A, Illumina b). We combined 500 µL of the denatured and diluted 16S rRNA library (1.8 pM) with 8 µL of denatured and diluted Illumina generated PhiX control library (1.8 pM) to assess sequencing error rates. For each run, we used four custom sequencing primers Read 1, Index 1, Index 2 and Read 2, which were diluted and loaded into the correct position of the reagent cartridge. An additional Index 2 sequencing primer was designed to enable the dual-index barcoding method on the MiniSeq (Pichler et al., 2018). Pooled libraries were sequenced on the Illumina MiniSeq platform at the GeoBio-Center LMU.

Demultiplexing and base calling were both performed using bcl2fastq Conversion Software v. 2.18 (Illumina, Inc.). We used USEARCH (Edgar, 2010 and 2013) and QIIME version 1.9.1 (Caporaso et al., 2010 and 2012) for MiniSeq read trimming and assembly, OTU picking and clustering at 97% sequence identity, which we previously tested with mock communities sequenced on the same platform (Pichler et al., 2018). The initial step was to assemble paired-end reads using the fastq_merge pairs command with default parameters allowing for a maximum of five mismatches in the overlapping region. Stringent quality filtering was carried out using the fastq_filter command. We discarded low quality reads by setting the maximum expected error threshold (E_max), which is the sum of the error probability provided by the Q score for each base, to 1. Reads were de-replicated and singletons discarded. OTU representative sequences were identified by BLASTn searches against the SILVA 16S rRNA SSU NR99 reference database release 132 (Quast et al., 2013). All operational taxonomic units (OTUs) assigned to Chloroflexi were aligned with SINA online v.1.2.11 (Pruesse et al., 2007) and inserted in a Maximum Likelihood RAxML phylogenetic tree selecting the best tree among 100 replicates, using ARB (Ludwig et al., 2004). Partial OTU sequences were added to the tree using the maximum parsimony algorithm without allowing changes of tree typology.

### Metagenomes and transcriptomes

Whole genome amplifications were performed on DNA extracts at dilution 10 times through a multiple displacement amplification (MDA) step of 6 to 7 hours using the REPLI-g Midi Kit (QIAGEN) and following the manufacturer’s instructions. MDA-amplified PCR products were then diluted to DNA concentrations of 0.2 ng μL^−1^ and used in metagenomic library preparations with the Nextera XT DNA Library Prep Kit (Illumina, San Diego), then quantified on an Agilent 2100 Bioanalyzer System (Agilent Genomics, Santa Clara) and normalized with the Select-a-Size DNA Clean and Concentrator MagBead Kit (Zymo Research, Irvine) as previously described (Vuillemin et al., 2019), diluted to 1 nM and pooled.

For site 15, total RNA extractions were obtained from 3.5 g of wet sediments using the FastRNA Pro Soil-Direct Kit (MP Biomedicals, Irvine) following the manufacturer’s instructions, with the addition of 4 μL glycogen (0.1 g × mL^−1^) to increase yield during precipitation of the RNA pellet, and final elution in 40 µL PCR-grade water (Roche). Extraction blanks were processed alongside to assess laboratory contamination. RNA extracts were quantified using the QuBit RNA HS Assay Kit (Thermo Fisher Scientific, Waltham). DNAse treatment, synthesis of complementary DNA and library construction were processed on the same day from 10 µL of RNA templates, without any prior MDA step, using the Trio RNA-Seq kit protocol (NuGEN Technologies, Redwood City). All libraries were quantified as described above, diluted to 1 nM and pooled for further sequencing on the MiniSeq platform (Illumina). For site 11 and 12, RNA yields were too low to achieve amplification and sequencing.

The MiniSeq reads were trimmed and paired-end reads assembled into contigs, and open reading frames (ORFs) extracted and functionally annotated as previously published (Ortega-Arbulú et al., 2019; Vuillemin et al., 2019; Orsi et al., 2020a). Paired-end reads were trimmed and assembled into contigs using CLC Genomics Workbench 9.5.4 (https://www.qiagenbioinformatics.com/), using a word size of 20, bubble size of 50, and a minimum contig length of 300 nucleotides. Reads were then mapped to the contigs using the following parameters (mismatch penalty = 3, insertion penalty = 3, deletion penalty = 3, minimum alignment length = 50% of read length, minimum percent identity = 95%). Coverage values were obtained from the number of reads mapped to a contig divided by its length (i.e. average coverage). Only contigs with an average coverage >5 were selected for ORF searches, and downstream analysis (Ortega-Arbulú et al., 2019). This protocol does not assemble ribosomal RNA (rRNA), and thus results are only discussed in terms of messenger RNA (mRNA).

Taxonomic identifications were integrated with the functional annotations, performing BLASTp searches of ORFs against an updated SEED (www.theseed.org) and NCBI RefSeq databases containing all predicted proteins from recently described high-quality draft genomes and single cell genomes from the NCBI protein database. Our database also included all fungal genomes from the NCBI RefSeq database (Orsi et al., 2018; Ortega-Arbulú et al., 2019). The total number of predicted proteins in the updated database was 37.8 million. We used the DIAMOND protein aligner version 0.9.24 (Buchfink et al., 2015). Cut-off values for assigning hits to specific taxa were performed at a minimum bit score of 50, minimum amino acid similarity of 60, and an alignment length of 50 residues. All scripts and code used to produce the analysis have been posted on GitHub (github.com/williamorsi/MetaProt-database), and we provide a link to the MetaProt on the GitHub page, as well as instructions within the scripts regarding how to conduct the workflows that we used.

We chose to focus on the coverage of total annotated protein-encoding ORFs detected, as opposed to the number of reads mapping per kilobase per ORF (for example, RPKM), to reduce potential bias from small numbers of “housekeeping” genes with potentially higher expression levels (Orsi et al., 2019 and 2020a). In addition, COG categories were assigned by comparing the metatranscriptome and metagenome annotated protein-encoding ORFs against the COG database (Galperin et al., 2015). Statistical analyses of beta-diversity were performed using RStudio v. 3.3.3 with the Bioconductor package (Huber et al., 2015). For both site 11 and 12, the metagenomes were sequenced for samples from four different depths to an average depth of 15 million reads, and *de novo* assembly resulted in a total of 177,498 contigs across all samples sequenced (Table S1). For site 15, metagenomes were sequenced for samples from five different depths at an average depth of 8.4 million reads (± 2.5 millions), assembled into 70,157 contigs. The metatranscriptomes from each sample at site 15 were produced in biological replicates and sequenced at an average depth of 4.0 million reads (± 1.5 million). *De novo* assembly resulted in a total of 91,199 contigs across all samples sequenced (Table S1). For all metagenomes and metatranscriptomes, the read coverage of annotated protein-encoding ORFs assigned to Chloroflexi was normalized to the coverage of all transcripts and results shown as % of read coverage.

Taxonomic assignment of protein-encoding genes to Chloroflexi among the SAR202 and Dehalococcoidia clades was further confirmed in our metagenomes by phylogenetic analysis of the RNA polymerase sigma factor (*RpoD*) gene proteins annotated as Chloroflexi, using alignments of 619 amino acid residues. Phylogenetic analyses of the predicted alpha and beta subunits of the *Dsr* and *apr* gene proteins were performed for all the corresponding annotated taxa in our metagenomes and metatranscriptomes, using 466, 433, 754 and 157 aligned amino acid sites respectively (Orsi et al., 2016). For each of the five marker gene phylogenies (*RpoD, DsrA, DsrB, aprA, aprB*), ORFs annotated to those genes from our annotation pipeline were aligned against their top two BLASTp hits in the NCBI-nr and SEED databases using MUSCLE (Edgar et al., 2004). Conserved regions of the alignments were selected in SeaView version 4.7 (Gouy et al., 2010), using Gblocks with the following settings: allowing for smaller final blocks, gap positions within the final blocks, and less strict flanking positions. Phylogenetic analysis of the resulting amino acid alignments of the predicted proteins were conducted in SeaView version 4.7 (Gouy et al., 2010), using RAxML (Stamatakis et al., 2012; Stamatakis, 2014) with BLOSUM62 as the evolutionary model and 100 bootstrap replicates. All alignments have been uploaded and will be made publicly available through the LMU Open Data website.

Data are publicly available through NCBI BioProject PRJNA473406 and PRJNA590088. Metagenomes from sites 11 and 12 have Short Read Archive (SRA) BioSample accession numbers SAMN10924458 and SAMN10924459, corresponding to run accession numbers SRX5372537 to SRX5372545. Metagenomes and transcriptomes from site 15 have accession number SAMN13317858 to SAMN13317870, corresponding to run accession numbers SRR10481880 to SRR1048192. The 16S rRNA data are available in SRA BioSample accessions SAMN10929403 to SAMN10929517 and SAMN13324854 to SAMN13324920.

## Acknowledgements

This work was supported primarily by the Deutsche Forschungsgemeinschaft (DFG) project OR 417/1-1 granted to W.D.O. The expedition was funded by the US National Science Foundation through grant NSF-OCE-1433150. Shipboard microbiology efforts were supported by the Center for Dark Energy Biosphere Investigations (C-DEBI grant NSF-OCE-0939564). This is a contribution of the Deep Carbon Observatory (DCO). It is C-DEBI publication XXX.

## Supporting Information

**Supplementary Figure S1.**
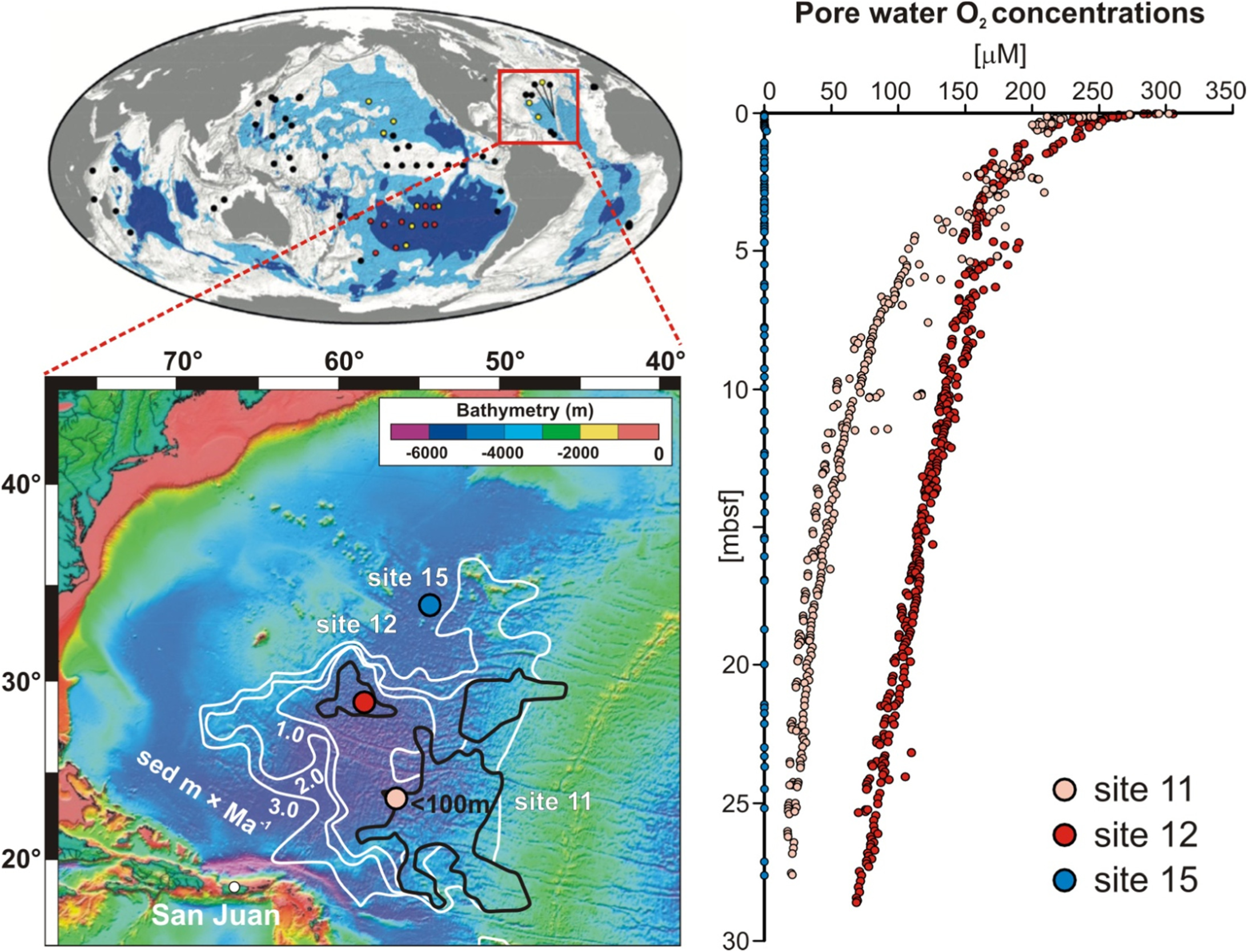
Map of sampling site 11, 12 and 15 with bathymetry, sedimentation rates and pore water O_2_ profiles. (**Left**) Global map of subseafloor sedimentation rates and corresponding O_2_ penetration is modified from D’Hondt *et al*. (2015). (**Right**) Note that at site 11 and 12, there is O_2_ penetrating to 30 mbsf, whereas at site 15 O_2_ is consumed immediately at the seafloor surface and the entire sediment sequence is anoxic.

**Supplementary Figure S2.**
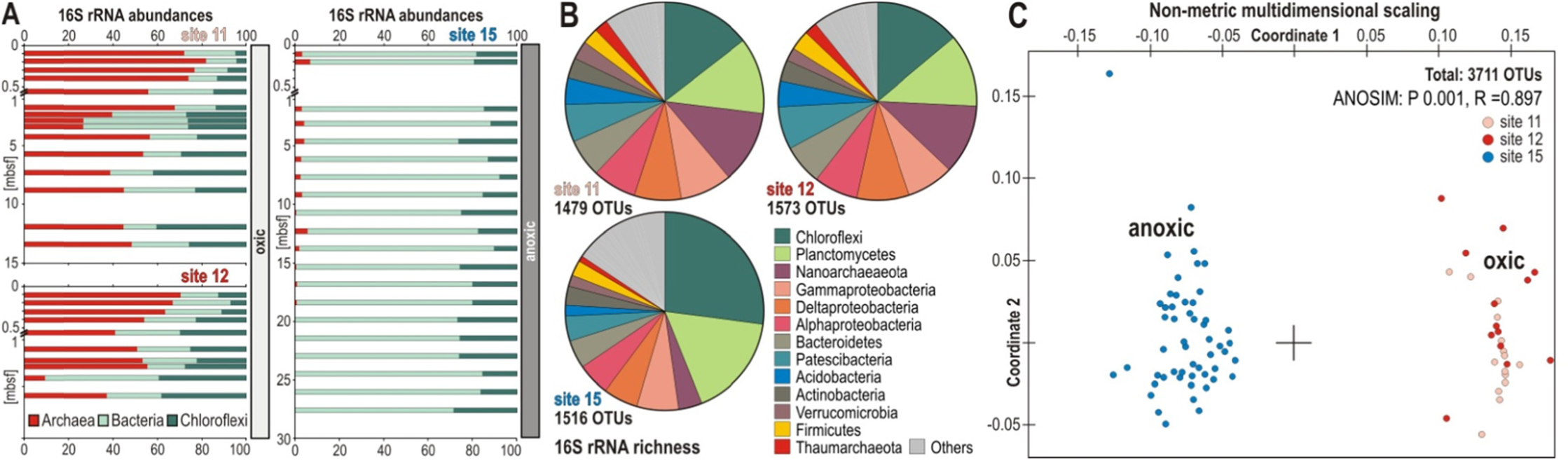
Microbial abundance, richness beta diversity based on 16S rRNA genes. (**A**) Downcore profiles for the relative abundances of Archaea (red), Chloroflexi (dark green) and all other Bacteria (light green). (**B**) Richness of 16S rRNA genes based on the total number of OTUs. (**C**) Non-metrical dimensional scaling (NMDS) plot for all three sites based on all OTUs obtained from 16S rRNA gene ampl**i**cons, with significant separation between oxic and anoxic sediment.

**Supplementary Figure S3:**
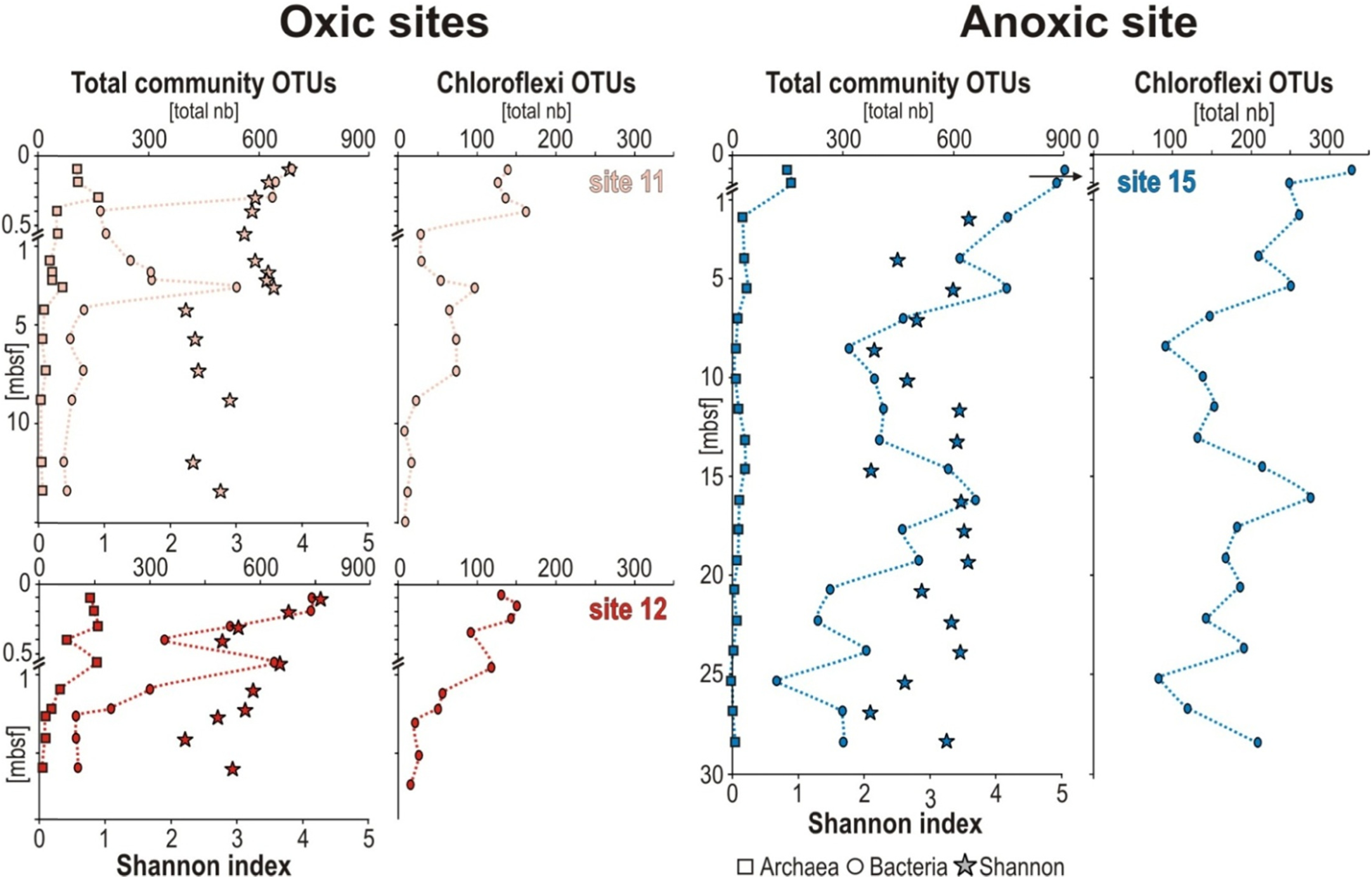
Alpha diversity for whole communities and Chloroflexi at the three sites. (**Left**) Total number of OTUs assigned to Archaea (squares), and Bacteria (circles) with Shannon indices (stars) calculated for the whole populations at the two oxic sites; number of OTUs assigned to Chloroflexi at the two oxic sites. (**Right**) Same indices calculated for populations from the anoxic site.

**Supplementary Figure S4:**
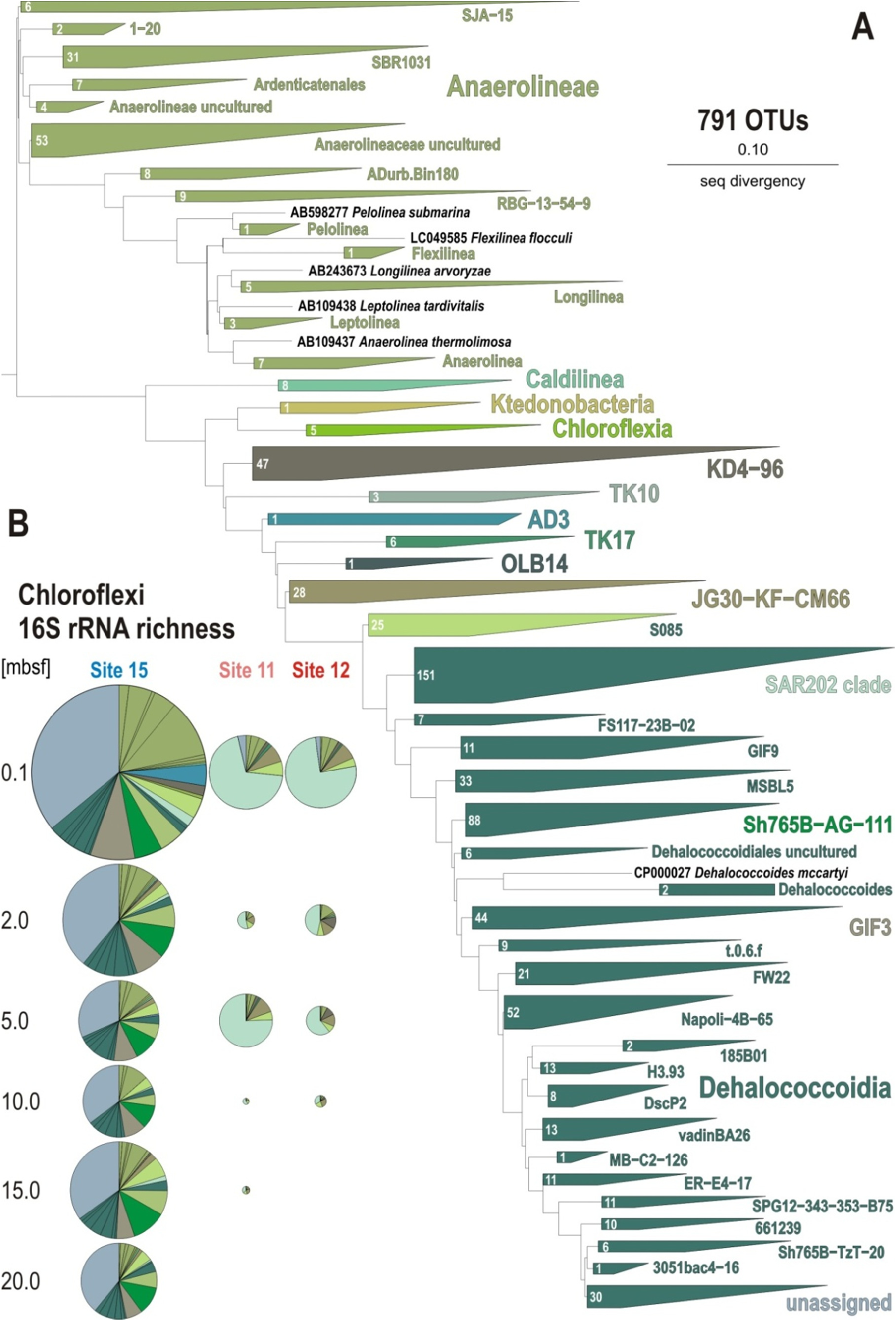
Phylogenetic analysis of 16S rRNA genes (V4 hypervariable region). (**A**) Phylogenetic tree obtained for all operational taxonomic units (OTUs) assigned to Chloroflexi in this study. (**B**) Pie charts displaying 16S rRNA gene richness of the whole assemblages of Chloroflexi for each of the three sites and at each sediment depth. The radii of pie charts are proportional to the number of OTUs.

**Supplementary Table S1:**
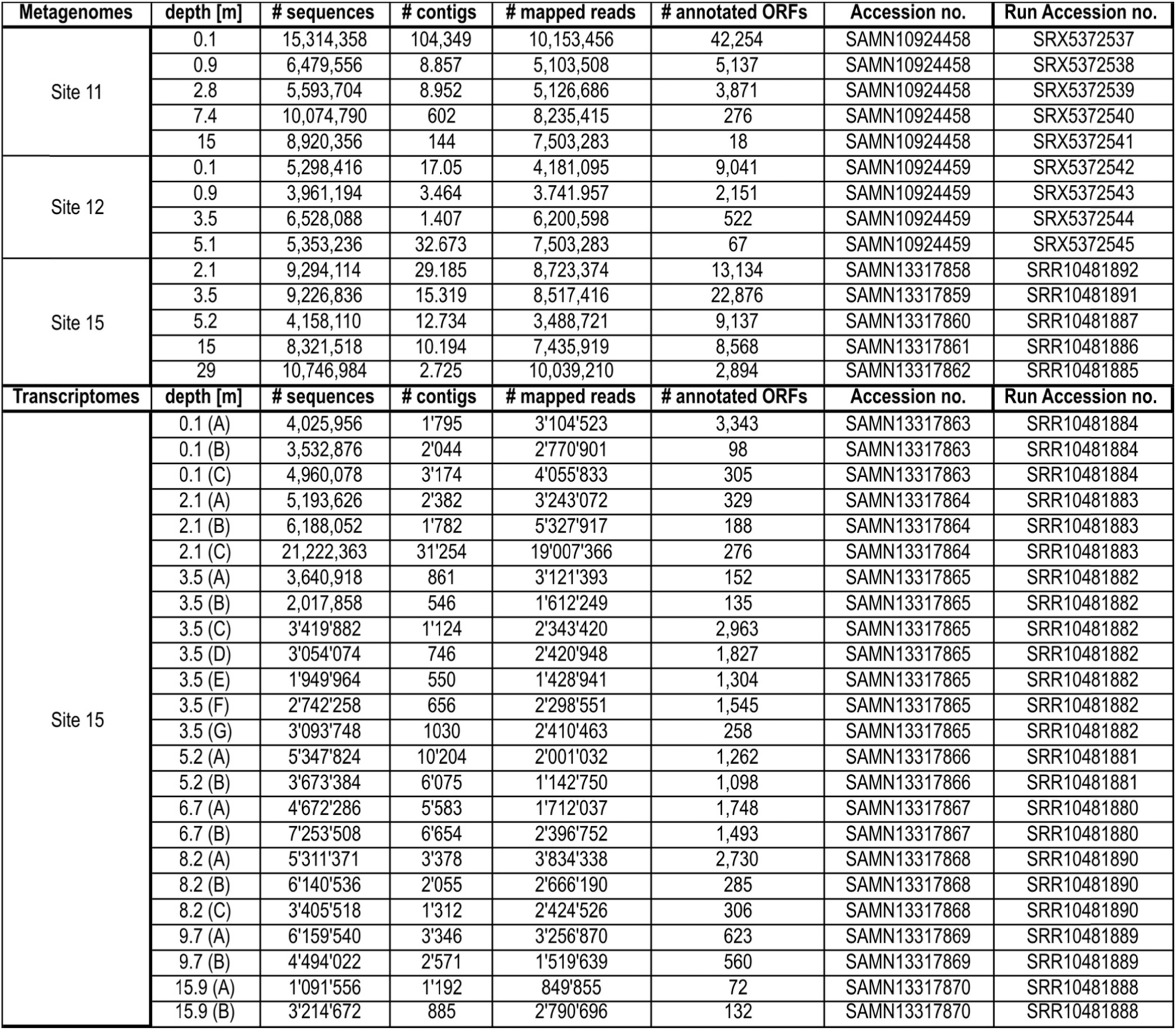
Sequencing and assembly results for the subseafloor metagenomes at site 11, 12 and 15, and transcriptomes at site 15 with their corresponding accession and run accession numbers.

